# Spatiotemporal dynamics of hydrogen peroxide during neutrophil swarming in 3D

**DOI:** 10.64898/2026.06.11.731567

**Authors:** Mathieu Deygas, Caroline Rogoll, Mathilde Bernard, Alexandre Deslys, Jeanne Lefévère-Laoide, Ivan Mikaelian, Moira Garcia-Gomez, Camille Plancke, Matthieu Piel, Pablo Vargas

## Abstract

Neutrophil swarming enables the coordinated recruitment of large numbers of cells to sites of infection. Although reactive oxygen species (ROS) are key mediators of neutrophil function, their dynamics during collective behavior remain poorly defined. Here, we developed a 3D ex vivo swarming system combined with HyPer7-expressing dHL-60 cells to visualize intracellular hydrogen peroxide (H_2_O_2_) dynamics in real time at single-cell resolution. This approach enabled us to reliably measure NADPH oxidase 2 (NOX2) activity during neutrophil swarming, revealing dynamic H_2_O_2_ production. We show that H_2_O_2_ production is spatially confined to cells at the stimulation site and temporally coupled to swarming initiation and distal cell recruitment. ROS production depends on NOX2 activity and calcium signaling but is not required for swarm formation or amplification. Notably, H_2_O_2_ accumulation was detected throughout activated cells rather than being restricted to phagosomes, suggesting a broader intracellular oxidative response. Together, these findings reveal a coordinated oxidative program associated with neutrophil swarm initiation and raise the possibility that ROS contribute to signaling functions beyond their established role in phagocytosis.

## INTRODUCTION

Neutrophils constitute the first line of defense against pathogens. They are rapidly recruited to sites of infection, where they deploy multiple mechanisms to eliminate their targets. Imaging studies have identified neutrophil swarming as a key innate immune strategy occurring in infected or inflamed tissues.^1,2^ During swarming, large numbers of neutrophils undergo coordinated chemotaxis to accumulate around and surround their target. This process relies on key molecular regulators controlling swarm initiation, amplification, and termination,^2–4^ including calcium-dependent production of leukotriene B4 (LTB4), which acts as a critical relay signal between neutrophils.^3,5–7^

Although neutrophil swarming has been observed in diverse inflammatory contexts in vivo,^2^ its study remains technically challenging due to the imaging requirements and the reliance on animal models. The development of on-chip swarming platforms has enabled major advances in the characterization of its molecular mechanisms.^6,8–11^ In these systems, neutrophils are exposed to micropatterned swarming stimuli, such as zymosan or heat-killed bacterial particles. However, these approaches do not recapitulate the physiological three-dimensional properties of tissue microenvironments. Despite recent progress in high-resolution imaging of swarm dynamics,^6,10,12,13^ a more comprehensive spatiotemporal understanding of both individual and collective neutrophil behavior during recruitment and within the swarm core remains needed, particularly in 3D tissue-like contexts.

Upon reaching their target, neutrophils deploy multiple antimicrobial mechanisms, including degranulation, neutrophil extracellular trap (NET) formation, and production of reactive oxygen species (ROS). ROS generation, known as the oxidative burst, constitutes a potent antimicrobial defense. Impaired ROS production, as observed in patients with chronic granulomatous disease (CGD), results in recurrent bacterial infections and tissue colonization.^14^ ROS-mediated antimicrobial activity can occur extracellularly or within phagosomes.^15,16^ Activated neutrophils generate superoxide via NADPH oxidase 2 (NOX2), which is converted into H_2_O_2_ and further gives rise to toxic oxygen derivatives, including hypochlorous acid produced by myeloperoxidase (MPO) within phagosomes. Beyond their antimicrobial role, ROS also regulate neutrophil functions by promoting granule release, NET formation, and inflammatory cytokine production.^17^ These responses must be tightly controlled to prevent excessive inflammation and autoimmune pathology.^18^

In the context of neutrophil swarming, it has been proposed that intra-versus extracellular ROS production contributes to scaling neutrophil recruitment according to target size.^15^ In addition, patients with CGD display dysregulated swarming with excessive neutrophil recruitment.^6,8,19^ Although this phenotype appears independent of ROS themselves,^6,20^ how NOX2 activation and ROS production are regulated during neutrophil swarming, and which cellular responses they influence, remain open questions.

Progress in addressing these questions has been limited by methodological constraints in monitoring ROS dynamics with specificity and spatial and temporal resolution. Existing approaches either lack specificity for distinct ROS species or fail to provide spatiotemporal resolution. For example, luminol-based chemiluminescence enables quantification of superoxide production but does not allow simultaneous imaging of cellular behavior.^15^ Chemical probes are often cumulative and poorly suited to capture the dynamic and reversible nature of ROS production.^12,21^ Recent advances have enabled improved spatiotemporal visualization of ROS.^21^ In particular, the genetically encoded probe HyPer7 allows highly sensitive, real-time, and specific detection of H_2_O_2_.^22^

Here, we used HyPer7-expressing neutrophil-like HL-60 cells (dHL-60) to characterize the spatiotemporal regulation of H_2_O_2_ production during neutrophil swarming in a 3D environment. We show that HyPer7 provides a robust readout of NADPH oxidase activation in this system. Using this approach, we found that H_2_O_2_ production is spatially restricted to cells located at the target site and is temporally coupled to swarm initiation. Intracellular H_2_O_2_ levels increased throughout activated cells, suggesting that ROS accumulation is not confined to phagosomes. Furthermore, H_2_O_2_ production depended on calcium signaling but was dispensable for swarm initiation and amplification. Together, these findings reveal a tightly coordinated oxidative response during neutrophil swarming and suggest that ROS may contribute to neutrophil functions beyond their established role in phagocytosis.

## RESULTS

### Real-time measurement of intracellular hydrogen peroxide in dHL-60 neutrophil-like cells

To monitor and visualize intracellular H_2_O_2_ levels in neutrophils, we generated an HL-60 cell line stably expressing the Hyper7 probe in all cellular compartments. Hyper7 is a fast, pH-stable, and ultrasensitive ratiometric probe for H_2_O_2_ (Figure 1A).^22^ Imaging of this probe relies on the acquisition of two channels: excitation at 475 nm for the oxidized form and at 405 nm for the reduced form (with identical emission at 525 nm), followed by calculation of the “HyPer7 ratio,” defined as the F475/F405 ratio (Figure 1B). Such ratiometric measurements eliminate biases caused by heterogeneous probe concentration or cell density. An increase or decrease in the Hyper7 ratio reflects a corresponding increase or decrease in average probe oxidation, respectively. This was confirmed when we monitored the response of differentiated HL-60 (dHL-60) neutrophil-like cells to exogenous H_2_O_2_ (Figure 1C). At a concentration of 20 µM, Hyper7 became rapidly oxidized, showing detectable signal increase already at the first acquisition time point (20 s after H_2_O_2_ addition). Subsequently, Hyper7 signal decreased, demonstrating the fast and reversible response of the probe under our imaging conditions. At higher concentrations of H_2_O_2_ (200 µM), HyPer7 oxidation reached a plateau that remained stable over the course of the recording, indicative of persistent ROS production (Figure 1C).

**Figure 1.**
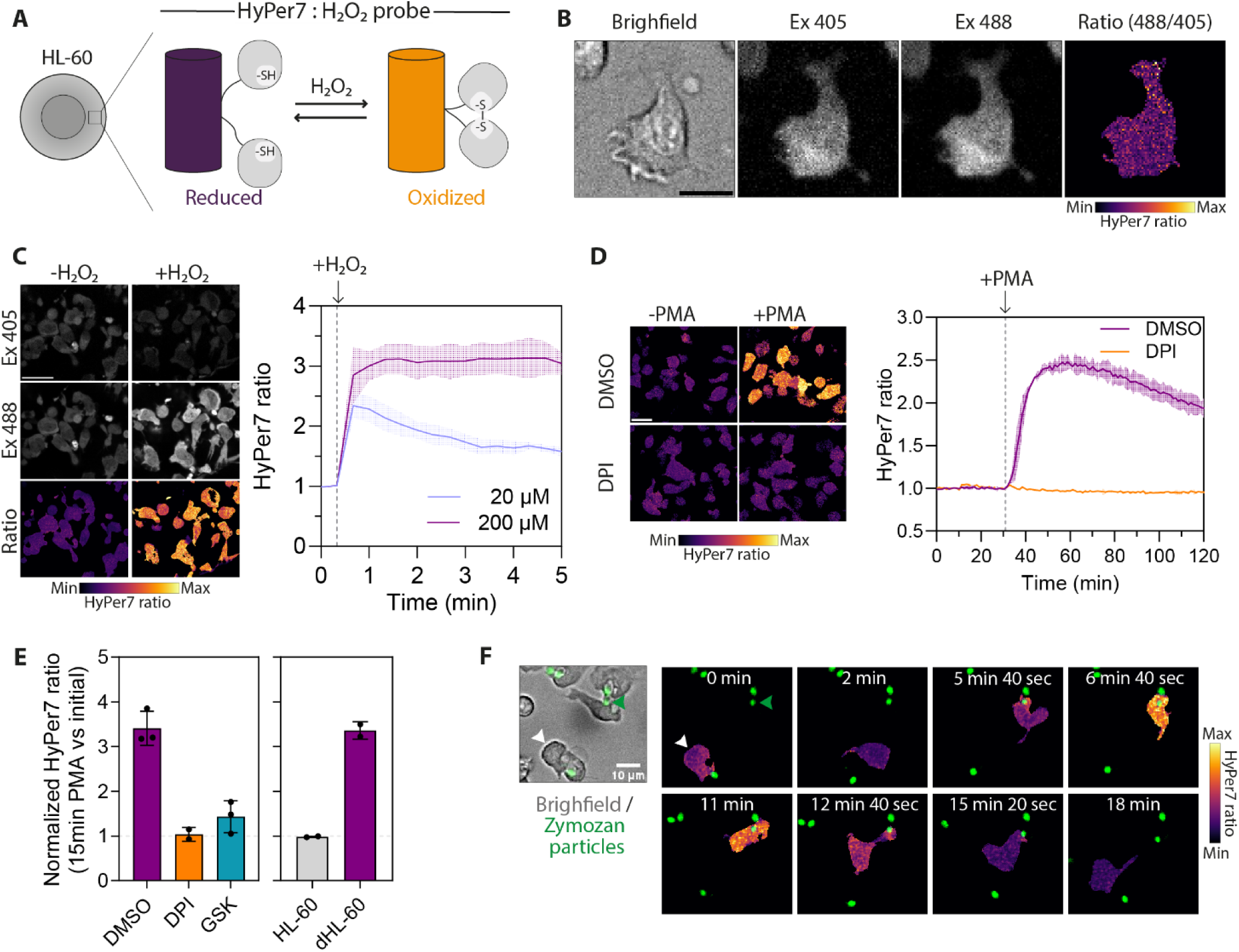
HL-60 HyPer7 cell line for studying spatiotemporal H_2_O_2_ production. (A) Schematic representation of the reduced and oxidized forms of the HyPer7 protein probe, stably expressed in HL-60 cells. (B) Imaging of the HyPer7 probe using excitation at 405 and 488 nm (with identical emission at 525 nm) and calculation of the HyPer7 ratio in a single HL-60 cell. Scale bar, 10 µm. (C) HyPer7 ratio in response to the addition of exogenous H_2_O_2_. Scale bar, 20 µm. (D) HyPer7 ratio in response to stimulation with 5 nM PMA, in the presence or absence of 10 µM DPI. Scale bar, 20 µm. (E) Normalized mean HyPer7 ratio measured 30 min after 100 nM PMA addition in dHL-60 treated or not with DPI or GSK2795039 (GSK), and in non-differentiated HL-60 cells. Mean +/− SD of n = 2 or 3 independent experiments. (F) Example of a transient increase in the HyPer7 ratio in a single neutrophil upon contact with a zymosan particle.

We next tested whether Hyper7 could also detect endogenous H_2_O_2_ generation. To this end, we used Phorbol 12-myristate 13-acetate (PMA), an activator of protein kinase C which can subsequently trigger NOX assembly. PMA stimulation triggered a rapid and transient intracellular H_2_O_2_ increase, peaking within 6-7 min before gradually declining over time (Figure 1D). Importantly, PMA-induced H_2_O_2_ production was abolished in the presence of the broad NOX inhibitor Diphenyleneiodonium (DPI), and upon specific NOX2 inhibitor (GSK2795039), confirming NOX2 as the source of H_2_O_2_ production (Figure 1D). Undifferentiated HL-60 cells, which lack NOX subunits expression, failed to produce H_2_O_2_ upon PMA stimulation (Figure 1E).^23^ In addition, contact of dHL-60 neutrophils with single zymosan particles induces a transient and global increase in intracellular H_2_O_2_ (Figure 1F).

Altogether, these results demonstrate that HyPer7-expressing dHL-60 cells constitute a robust tool to monitor NOX2 activation and intracellular H_2_O_2_ dynamics in neutrophil-like cells. This probe enables single-cell resolution measurements of H_2_O_2_ production on a timescale of seconds and captures its dynamics in a reversible manner.

### Establishment of a 3D in vitro system to study dHL-60 neutrophil swarming

Existing in vitro swarming assays usually rely on unconfined systems in which weakly adherent neutrophils migrate towards swarming stimuli in the absence of physical challenge. To investigate swarming in a more physiologically relevant microenvironment, we developed a simplified in vitro system in which cells swarm as they are embedded within a 3D collagen matrix. Briefly, zymosan spots were manually deposited onto a glass-bottom microwells (Figure 2A). This approach generates zymosan spots of variable sizes, unlike previously described tightly controlled systems.^8,9,24^ (Figure 2A). Upon zymosan spot drying, a low-density collagen gel (1 mg/ml) containing dHL-60 neutrophils was cast on top. The low density allows a substantial fraction of cells to sediment near the glass surface during collagen polymerization, allowing cells accumulation close to the glass and zymosan spots while preserving a 3D microenvironment (Figure 2A). To facilitate cell tracking during swarming, we mixed cell populations labelled with distinct nuclear dyes (Figure 2B). Live cell imaging was initiated immediately after collagen polymerization (10 min), enabling continuous tracking of cell migration along swarming formation.

**Figure 2.**
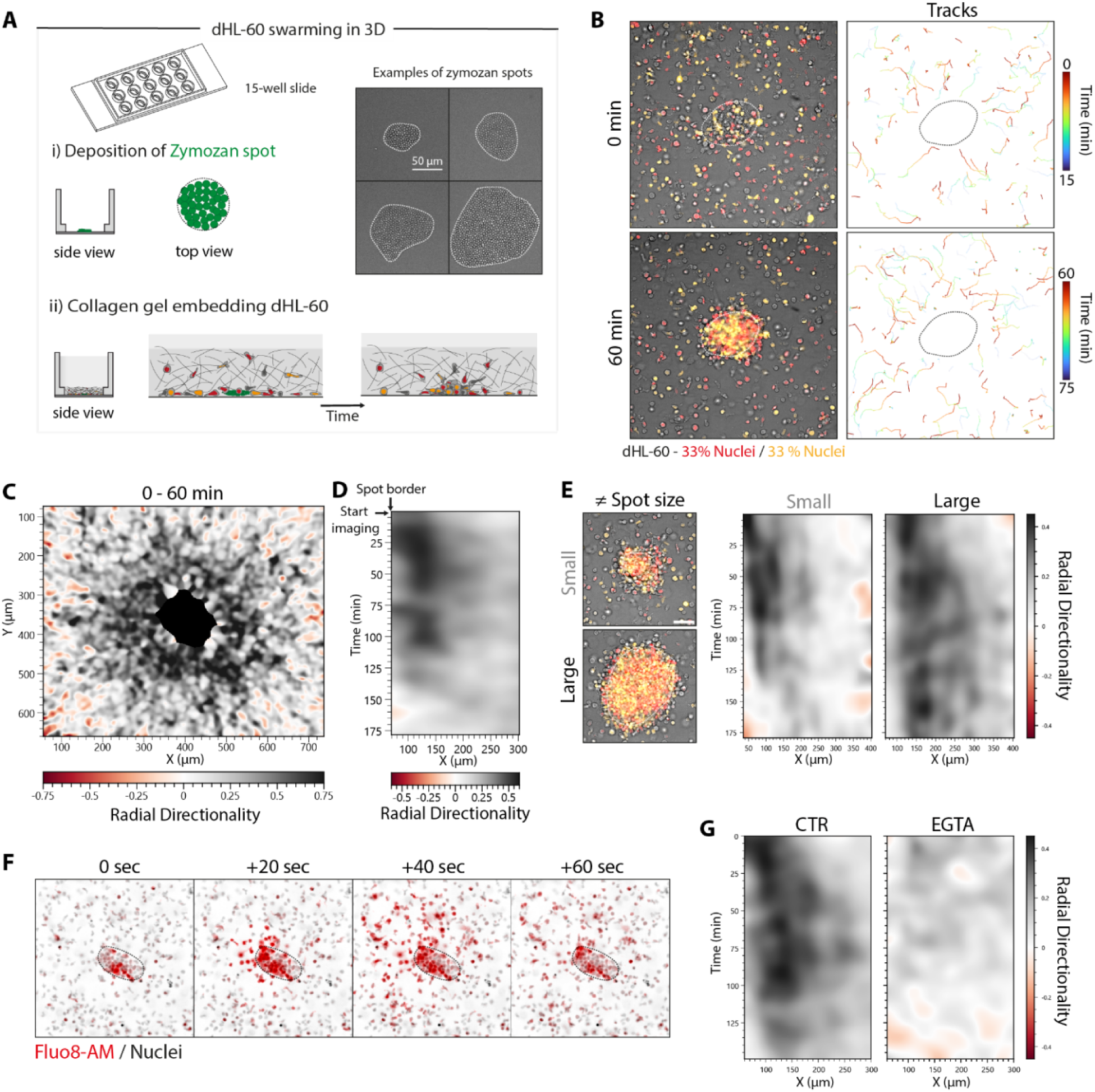
dHL-60 swarming using in vitro 3D system. (A) Schematic representation of the in vitro system used to study neutrophil swarming in 3D. A zymosan spot is manually deposited onto a glass surface (i), and a collagen gel containing differentiated HL-60 (dHL-60) cells is cast on top (ii). The well is then placed at 37 °C to initiate imaging. (B) Imaging and tracking of dHL-60 using a mix population composed of differentially stained nucleus. The dotted line represents the zymosan spot boundary. (C) Contour plot (XY) showing mean radial directionality between 0 and 60 min of imaging. N = 4 swarms from one representative experiment. (D) Contour plot (XT) showing mean radial directionality evolution over time, from (C). (E) Contour plot (XT) showing mean radial directionality evolution over time for small or large spots. N = 3 swarms from one representative experiment. (F) Time-lapse sequence of a multicellular wave of neutrophil calcium activity following detection of the zymosan. (G) Contour plot (XT) showing mean radial directionality evolution over time for control and EGTA treated cells. N = 5 swarms from one representative experiment.

In 3D, dHL-60 cells exhibited robust swarming behavior, characterized by a progressive accumulation of cells at the zymosan spot over time (Figure 2B). Detailed cell trajectory analysis of dHL-60 swarming towards zymosan spots (diameter 120–150 µm) revealed strong radial directionality toward the center of the swarm within a ∼250 µm radius from the edge of the spot. To avoid tracking errors due to high cell density, dHL-60 within the spot area itself were excluded from the analysis (black region at the center) (Figure 2C). Spatiotemporal analysis showed that directed neutrophil recruitment persisted for approximately 2 hours before progressively declining toward nondirectional migration (Figure 2D). These data show fast and persistent swarming of dHL-60 cells toward zymosan in 3D.

We next compared swarming dynamics in function of the size of the zymosan spots. Larger spots (diameter ∼200 µm) induced a broader recruitment zone, which reached about 300 µm from the zymosan edge. In contrast, smaller spots (diameter ∼50 µm) resulted in a lower recruitment zone of about 150µm (Figure 2E). As a main difference, and despite similar initial cell numbers, larger spots led to sustained cell recruitment for at least 3h, which was double as compared with smaller zymosan spots (Figure 2E and Figure S2E). These observations are consistent with previous findings showing that human neutrophils accumulate at targets proportionally to target size.^13,25^

We next asked whether this 3D system could also recapitulate intracellular calcium dynamics, a key regulator of neutrophil swarming. Using the calcium probe Fluo8-AM, we observed calcium activity waves initiating in cells located at the spot and propagating outward toward surrounding neutrophils (Figure 2F), as previously reported in unconfined swarming.^5–7^ Similar results were obtained using dHL-60 cells stably expressing the calcium sensor GCaMP6F (Figure S1). In the presence of EGTA, an extracellular calcium chelator, both calcium waves and calcium activity within cells at the swarm core were abolished (Figure S1A and S1B). In parallel, swarm initiation was completely inhibited upon calcium chelation (Figure 2G). These results show that 3D swarming reproduces key requirements previously described in confinement-free systems.

To assess the differences with an unconfined context, we next compared dHL-60 swarming freely in 2D versus in a confined 3D context. We found markedly reduced radial directionality in 2D relative to the 3D system (Figure S2A to S2D). Cells also responded over a smaller radius from the spot edge in 2D (Figure S2B and S2C). In addition, in 2D we observed frequent flow artifacts, thereby preventing efficient swarming and limiting quantitative analysis (Figure SA). Altogether, these results highlight that this simple system provides a robust and tractable platform for studying swarming and ROS dynamics in 3D matrices in vitro.

### Spatiotemporal H_2_O_2_ production during neutrophil swarming

To investigate the spatiotemporal dynamics of ROS production during neutrophil swarming, we used HyPer7-expressing dHL-60 cells in our 3D swarming system. Confocal live-cell spinning-disk imaging showed swarming initiation by cells directly contacting the zymosan spot within minutes. Monitoring HyPer7 probe signal revealed a marked increase in H_2_O_2_ production in cells located at the zymosan spot (Figure 3A and 3B). This increase was spatially restricted, as radial analysis showed a sharp decline in H_2_O_2_ levels at the edge of the zymosan spot, reaching basal levels beyond this boundary (Figure 3A and 3C). For quantitative analysis, we defined two regions: an “inside” region corresponding to the zymosan spot and an “outside” region containing cells located beyond. H_2_O_2_ levels in the outside region remained at basal levels throughout the recording period. In contrast, H_2_O_2_ levels within the inside region progressively increased, reached a maximal average value after ∼30 min, and remained elevated for the duration of the acquisition (80 min) (Figure 3D and S3A). However, the temporal dynamics of H_2_O_2_ production varied across swarms. Two main profiles were observed: i) a sustained response, in which H_2_O_2_ levels plateaued after reaching a maximum; and ii) a transient response, characterized by a decline following the peak (Figure 3E). This heterogeneity appeared to be associated with differences in zymosan spot size. Indeed, analysis of swarms of varying sizes revealed that smaller swarms displayed transient H_2_O_2_ responses, whereas larger swarms exhibited more sustained H_2_O_2_ production (Figure 3F and 3G). In smaller spots, the decline in H_2_O_2_ levels after peak response was associated with degradation and dispersion of zymosan particles from their original location (Figure S3B). Notably, the amplitude of H_2_O_2_ increase negatively correlated with zymosan spot size (Figure S3C).

**Figure 3.**
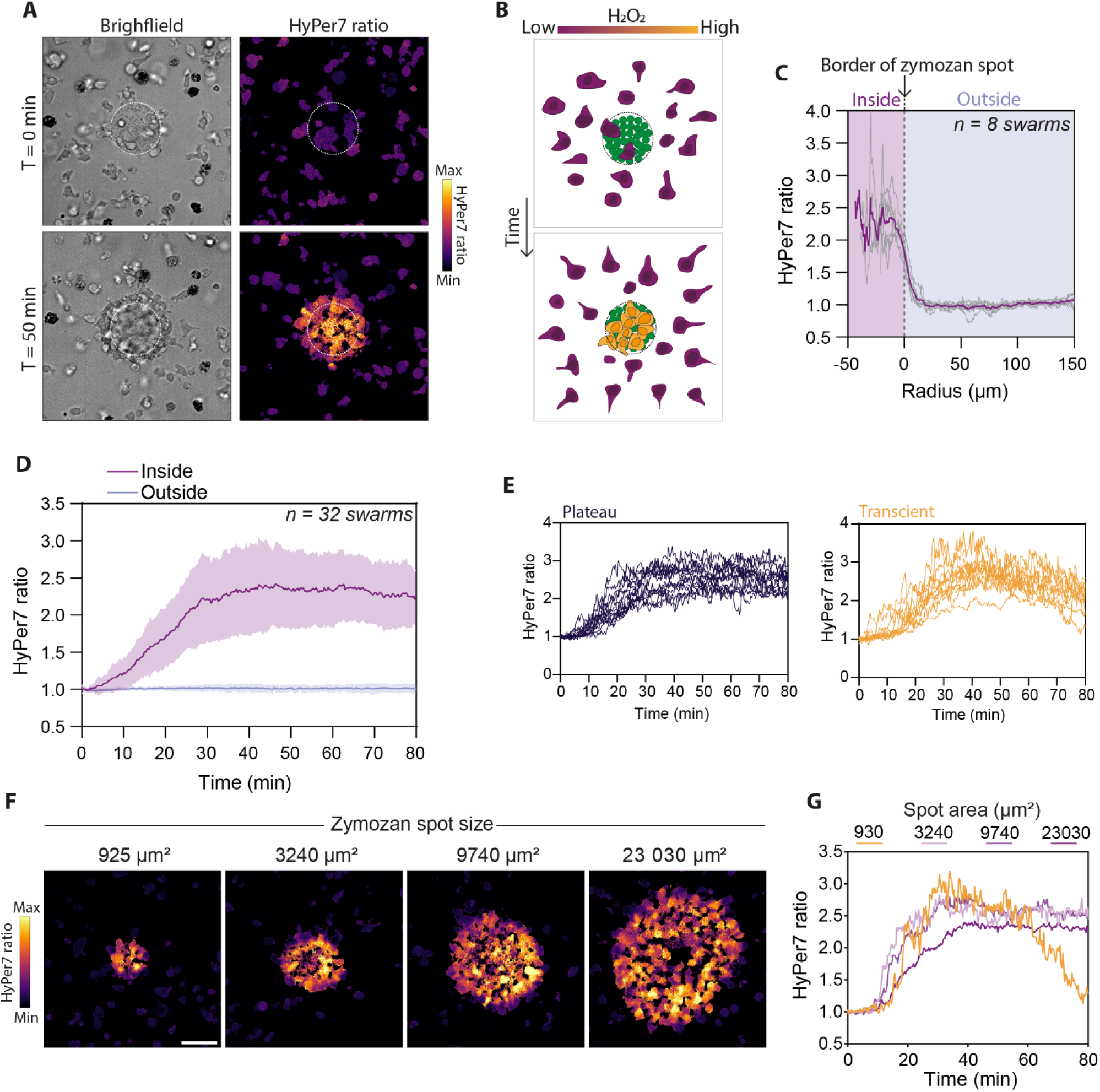
Spatiotemporal analysis of H_2_O_2_ levels during dHL-60 swarming. (A) HyPer7 ratio of dHL-60 before swarming initiation (T = 0 min) and during active swarming (T = 50 min) using the 3D system. The dotted line represents the zymosan spot boundary. (B) Schematic representation of neutrophil swarming and the associated increase in H_2_O_2_ levels specifically in cells within the swarm. (C) Radial analysis of H_2_O_2_ levels demonstrating that the increase is restricted to cells within the swarm. N = 8 swarms from 3 independent experiments. (D) Spatiotemporal analysis of H_2_O_2_ levels inside and outside the initial zymosan area, during the first 80 min of imaging. N = 32 swarms from 5 independent experiments. (E) Temporal evolution of mean intracellular H_2_O_2_ levels (inside region) across swarms, categorized into two behavioral patterns: plateau and transient. (F) Median intracellular H_2_O_2_ levels measured over a 10-min window (centered at t = 30 min) for four swarms of different sizes. (G) Temporal evolution of mean intracellular H_2_O_2_ levels (inside region) for the four swarms shown in (G).

Together, these results demonstrate that neutrophils rapidly produce H_2_O_2_ during the early stages of swarming. This response remains spatially confined to cells located within the zymosan region, revealing a localized oxidative response associated with collective neutrophil recruitment.

### Coordinated and calcium-dependent H_2_O_2_ production is tightly coupled to swarming initiation

We next examined the temporal relationship between H_2_O_2_ production and neutrophil swarming. In our system, swarming initiation occurred after a variable delay (Figure S2A), likely reflecting differences in the number of cells initially present at the zymosan spot and/or their ability to trigger swarm amplification. Interestingly, the increase in H_2_O_2_ levels coincided with swarm initiation and with the recruitment of cells located away from the zymosan spot (Figure 4A and 4B). Consistently, the onset of H_2_O_2_ elevation strongly correlated with the initiation of distant cell recruitment, with a slope close to 1 (Figure 4C and 4D). In rare cases, swarming failed to initiate despite the presence of cells at the zymosan spot (Figure S4A). Under these conditions, no global increase in H_2_O_2_ was observed. Instead, individual cells displayed transient H_2_O_2_ production, similar to the response observed when a single cell encounters a zymosan particle (Figure 1E). Furthermore, in the presence of EGTA, which prevents swarm initiation (Figure 2G), the increase in H_2_O_2_ was completely abolished (Figure 4E). Together, these observations indicate that global H_2_O_2_ production is tightly associated with, and dependent on, swarm initiation.

**Figure 4.**
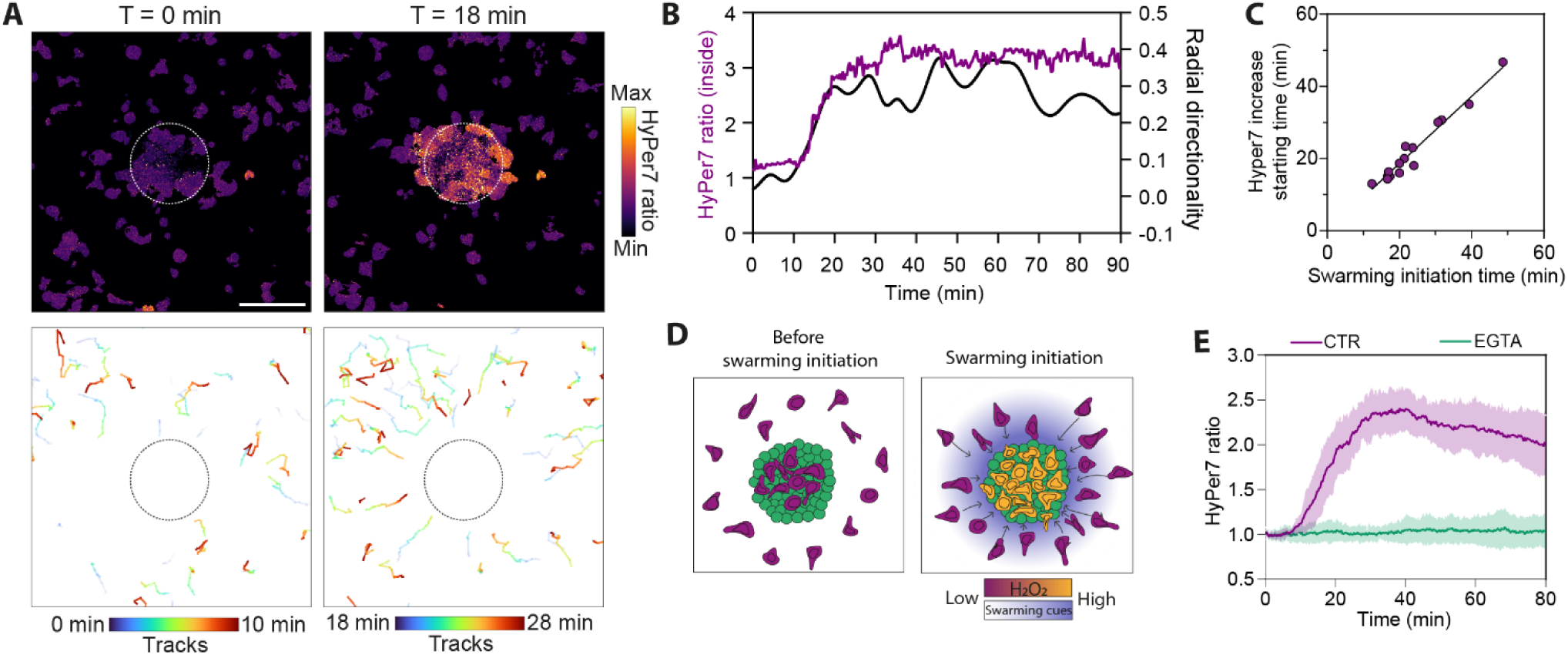
H_2_O_2_ increase coincides with swarming initiation. (A) HyPer7 ratio and tracking of dHL-60 before swarming initiation (T = 0 min) and at swarming initiation (T = 18 min) using the 3D system. The dotted line represents the zymosan spot boundary. Scale bar, 50 µm. (B) Evolution of mean HyPer7 ratio (inside the zymosan area) and radial directionality (of cells located outside) over time, from the swarm represented in (A). (C) Correlation between the starting time of HyPer7 ratio increase and swarming initiation time. N = 16 swarms from n = 3 independent experiments. (D) Schematic illustrating the temporal correlation between swarming initiation and H_2_O_2_ elevation. (E) Spatiotemporal analysis of H_2_O_2_ levels within the zymosan area in the presence or absence of 2 mM EGTA. N = 10 swarms from 2 independent experiments.

While H_2_O_2_ production appeared predominantly as a coordinated global response, we observed some heterogeneity in larger swarms. In these conditions, localized and transient H_2_O_2_ signals were detected in subgroups of cells within discrete regions (Figure S4B), suggesting that ROS production can also arise from local signaling events. Of note, H_2_O_2_ elevation was not associated with cell death, which occurred only at later stages of the experiment, starting approximately 3 h after swarm initiation (Figure S5A and S5B). Because the HyPer7 probe freely diffuses throughout the cytoplasm, it does not allow precise subcellular localization of H_2_O_2_ production. Interestingly, nuclear-targeted HyPer7 also revealed increased H_2_O_2_ levels, consistent with a global increase in intracellular ROS within activated cells (Figure S5C).

Together, these results indicate that H_2_O_2_ production is a coordinated and temporally synchronized response among cells at the zymosan spot, tightly coupled to swarming initiation and dependent on calcium signaling. These findings suggest that ROS production is not solely a cell-autonomous consequence of zymosan uptake, but is instead integrated into the collective dynamics of the swarming program

### NOX2-dependent H_2_O_2_ production is dispensable for swarming initiation and amplification

Given the relationship between swarming initiation and H_2_O_2_ elevation, we next asked whether inhibition of H_2_O_2_ production affects swarming dynamics. As expected, treatment with either the pan-NOX inhibitor DPI or the NOX2-specific inhibitor GSK2795039 abolished H_2_O_2_ production (Figures S6A and S6B), confirming NOX2 as the source of H_2_O_2_ within the swarm. In the presence of either inhibitor, swarming initiation was not affected (Figures 5A and 5B). With DPI treatment, we observed a trend toward a faster loss of radial directionality compared with control conditions. However, this effect was less pronounced with the NOX2-specific inhibitor, suggesting a possible off-target effect of DPI. Together, these results indicate that H_2_O_2_ production is not required for either swarming initiation or swarm amplification.

**Figure 5.**
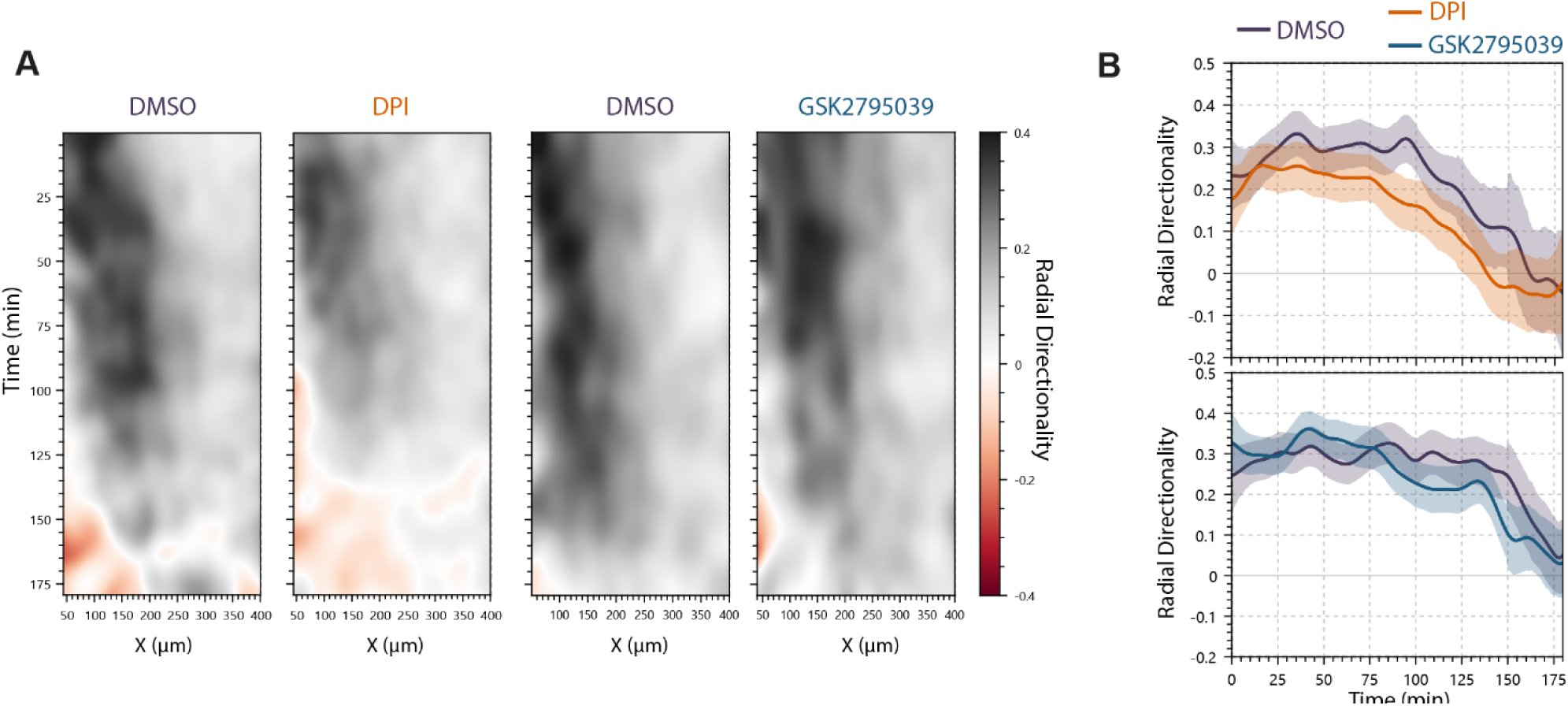
NOX2-dependent H_2_O_2_ production is dispensable for swarming dynamic. (A) Contour plot (XT) showing mean radial directionality evolution as a function of time for DMSO, DPI, and NOX2i treated cells. N = 10 swarms (DMSO vs DPI) and 8 swarms (DMSO vs GSK2795039) from 3 independent experiments. (B) Evolution radial directionality as a function of time, in the selected radius of 200 µm from (A).

## DISCUSSION

Neutrophil swarming is a hallmark of innate immune responses, yet the spatiotemporal regulation of reactive oxygen species during this collective behavior remains poorly understood. In this study, we combined a 3D ex vivo swarming model with the genetically encoded H_2_O_2_ sensor HyPer7 to visualize intracellular ROS dynamics at single-cell resolution. Using this approach, we show that H_2_O_2_ production is rapidly induced during swarming, remains spatially restricted to cells located at the target site, and is temporally coupled to swarm initiation. We further demonstrate that this response depends on NOX2 activity and calcium signaling, yet is dispensable for swarm initiation and amplification. Together, these findings reveal a coordinated oxidative response associated with the onset of neutrophil swarming and establish a framework for investigating how ROS contribute to neutrophil functions during collective immune responses.

Previous study reported that HL-60 cell swarming is less robust and less organized than swarming by primary human neutrophils, resulting in smaller and less coordinated swarms.^24^ By embedding cells within a collagen matrix, our 3D swarming system markedly improves the robustness of dHL-60 swarming behavior. In contrast to conventional 2D systems, where cell movements are frequently affected by fluid flow, the collagen matrix physically constrains cells and limits passive displacement, thereby improving the reliability of cell tracking and behavioral analyses. The 3D environment may also reduce the diffusion of signaling molecules, facilitating their detection by neighboring cells and promoting signal amplification and recruitment within the swarm. In addition, local remodeling or degradation of the collagen matrix by clustered cells could generate chemoattractant peptides,^26^ further enhancing neutrophil recruitment. Together, these features establish this simple and physiologically relevant 3D system as a powerful ex vivo platform for studying neutrophil swarming.

The study of neutrophil ROS production remains technically challenging. Current approaches are often limited by the lack of specificity for individual ROS species, the inability to simultaneously visualize cells, as in luminol-based chemiluminescence assays,^15,27^ or the absence of temporal resolution when using cumulative chemical probes such as DCFH-DA or BioTracker Orange OH and HClO Dye.^12,28^ The genetically encoded HyPer7 probe overcomes many of these limitations and enables specific visualization of intracellular H_2_O_2_ dynamics in living cells. Using HyPer7-expressing HL-60 cells, we show that H_2_O_2_ production is spatially restricted to cells located within the zymosan spot region. These observations are consistent with previous findings obtained using chemical probes detecting OH and HClO, ROS species generated downstream of H_2_O_2_ and associated with phagocytosis.^12^ Our work further provides new insights into the temporal dynamics of ROS production. In particular, H_2_O_2_ elevation was transient in small swarms, declining within minutes after initiation. Moreover, the increase in H_2_O_2_ coincided temporally with the onset of neutrophil recruitment, suggesting that signals involved in swarm initiation may also contribute to NOX2 activation and ROS production. Consistently, H_2_O_2_ production was abolished in the presence of EGTA, which also prevents swarm initiation. This observation is notable because NOX2 itself does not contain EF-hand calcium-binding domains and is not considered to be directly calcium-dependent.^29^

Several questions nevertheless remain unresolved. First, why is ROS production restricted to cells located at the zymosan spot? One possible explanation is that NOX2 activation requires the integration of two distinct signals: a swarming-independent signal provided by direct contact with zymosan and a second signal released by neighboring cells during swarm initiation. Another important question concerns the precise localization of ROS production. Neutrophils can generate ROS both intracellularly and extracellularly.^15^ However, in our system, the HyPer7 probe does not allow discrimination between intracellularly generated H_2_O_2_ and extracellular ROS that subsequently diffuse into the cell. In addition, neutrophil ROS production may remain confined to phagosomes or instead occur at other intracellular sites, including the plasma membrane or mitochondria.^27,30–32^ Recent studies using HyPer7 have shown that H_2_O_2_ can be generated as localized hotspots with limited intracellular diffusion.^33,34^ At our imaging resolution, we primarily observed a global intracellular increase in H_2_O_2_ levels. Because HyPer7 freely diffuses throughout the cytoplasm, it is unlikely to resolve such local hotspots. Nevertheless, targeting HyPer7 specifically to the nucleus also revealed increased H_2_O_2_ levels during the early stages of swarming, although with lower amplitude than the cytosolic probe. These observations suggest that H_2_O_2_ production increase globally within neutrophils during swarming and are not exclusively restricted to phagosomes or to a single intracellular compartment.

In this context, what could be the functional consequences of the global intracellular increase in H_2_O_2_ observed during swarming? Beyond their well-established antimicrobial activity, ROS, and H_2_O_2_ in particular, owing to its relative stability, are increasingly recognized as signaling molecules. For example, in zebrafish, tissue injury induces H_2_O_2_ production by epithelial cells, leading to lipid peroxidation and activation of the Src-family kinase Lyn, thereby promoting neutrophil recruitment to the wound site.^35–37^ ROS produced by neutrophils themselves, in response to chemoattractant, can also modulate their migratory behavior. NADPH oxidase-derived ROS negatively regulate actin polymerization through reversible actin glutathionylation.^38^ In addition, fMLP-induced ROS production promotes oxidation of the ion channel TRPM2, leading to internalization of the fMLP receptor FPR1 and termination of neutrophil migration.^39^ Given the importance of receptor desensitization and stop signals in arresting neutrophil migration within the swarm core,^10^ the coordinated increase in H_2_O_2_ observed during swarming could contribute to these processes by limiting migratory responses once cells reach the target. ROS may also influence neutrophil effector functions at the target site. NOX2-dependent ROS generation has been implicated in NET formation,^40,41^ raising the possibility that the coordinated oxidative burst contributes to NET release observed during swarming.^8,42^ Furthermore, the rapid consumption of oxygen associated with oxidative burst activity may locally generate hypoxic microenvironments, as reported in inflamed tissues in vivo,^43^ potentially creating conditions that limit bacterial dissemination.

### Limitations of the study

An important limitation of this study is that we did not identify the functional role of the observed H_2_O_2_ increase or its potential impact on neutrophil functions beyond the well-established role of ROS production during phagocytosis. In addition, all experiments were performed using dHL-60 cells, and validation in primary human neutrophils will therefore be required. Moreover, zymosan was the only swarming stimulus used in this study. It will be important to determine whether similar ROS dynamics are observed in response to other bacterial or fungal stimuli. Previous studies have shown that both the magnitude and the source of neutrophil ROS production can vary substantially depending on the pathogen encountered.^27,44^ Finally, our observations were obtained exclusively in vitro. It will therefore be important to investigate whether similar H_2_O_2_ dynamics occur in vivo, where tissue oxygen levels are substantially lower than under standard in vitro conditions.

## MATERIAL AND METHODS

### Cell culture and differentiation

The human myeloid leukemia-derived promyelocytic cell line HL-60 was a kind gift of F.Sepulveda (Institut Imagine, Paris, France) and maintained in Roswell Park Memorial Institute (RPMI) 1640 Medium (Thermo Fisher Scientific, 11875093) supplemented with 10 % heat-inactivated fetal bovine serum (Biowest, S1810), 20 mM Hepes (Gibco, 15630056), and 1 % Penicillin-Streptomycin (Thermo Fisher, 15140-122). Cultures were maintained in a humified incubator at 37°C and 5 % CO_2_. Cells were passaged every 2-3 days to maintain a density between 0.2 and 1.0 x 10^6^ cells/mL and kept in polystyrene T25 flasks. For differentiation into a neutrophil-like cells (dHL-60), cells were cultured for up to 6 days in standard culture medium supplemented with 1.3 % dimethyl sulfoxide (DMSO). The medium was changed every 2-3 days during differentiation period. For all experiments, dHL-60 cells were used between day 5 and 6 post-differentiation. For imaging media (swarming and HyPer7 imaging), 5 % FBS supplemented media was used.

Human embryonic kidney (HEK) 293T cells were a kind gift of I.Mikaelian (CRCL, Lyon, France) and were cultured in Dulbecco’s modified Eagle’s medium (DMEM) supplemented with 10% FBS and 1 % Penicillin-Streptomycin. 293T cells were used to generate lentiviral particles HyPer7, NLS-HyPer7 and GCaMP6F probes in HL60 cell lines.

### HL-60 transduction

To obtain viral particles, HEK293T cells were cultured in DMEM supplemented with 10 % FBS, were used to generate lentiviral particles for HyPer7, NLS-HyPer7 or GCaMP6F expression in HL-60 cell line. To obtain lentiviral vector of interest, genes of interest were amplified from pCS2+HyPer7 (#136466 addgene), or pCS2+HyPer7-NLS (#136468 addgene) or pGP-CMV-GCaMP6f (#40755 addgene) and were cloned into pCSII-EF-MCS backbone. Lentiviral packaging plasmids pMD2.G (0.3 µg) and psPax2 (0.8 µg), along with 0.9 µg of lentiviral vector were transfected to 1.6 x 10^6^ HEK293T in a well of 6-well plate using Lipofectamine 2000 transfection reagent. After 24h, media was changed by HL-60 culture media. The culture supernatant containing lentiviral particles was collected after 48 h. Lentiviral particles supernatants were filtered with 0.45 µm filter, mixed gently with protamine sulfate 10 µg/mL and added to 1.5 x 10^5^ HL-60 cells for spinfection (1000 g for 2 h at 35°C). Cells were resuspended in complete media and sorted by flow cytometry to select GFP+ cells.

### Biochip preparation for 3D swarming assays

3D zymosan swarming chips were prepared using either 15-well µ-Slides (IBIDI, Cat.No:81506) or custom-made 35-mm FluoroDishes. For the latter, a polydimethylsiloxane (PDMS) chip containing 4-mm diameter wells was bonded to the glass bottom of the dish by plasma treatment. Zymosan patterns were manually deposited onto both types of substrates using a stamping procedure. A stock suspension of unlabeled zymosan particles (Zymosan A from S. cerevisiae, Thermo Fisher Scientific, Z2849) was prepared in PBS at a concentration of 20 mg/mL and supplemented with 2 mM sodium azide. We validated our main results with zymosan particles suspension stored without sodium azide. Custom stamps were fabricated by punching cylindrical PDMS pieces (0.5 mm diameter) from a spare PDMS slab using a biopsy punch. The stamp was held with tweezers, briefly dipped into a vortexed zymosan suspension, and manually pressed onto the substrate to generate five zymosan spots per well. Spot quality and morphology were verified by microscopy before use. Devices containing dried zymosan spots were prepared either on the day of the experiment or several days in advance and stored at 4°C until use.

### Collagen gel preparation

For swarming experiments, dHL-60 cells were embedded in a 1 mg/mL collagen gel. All reagents were kept on ice throughout the preparation procedure. A cell suspension was prepared at a concentration of 35 × 10⁶ cells/mL in imaging medium. The collagen mixture was prepared by sequentially combining imaging medium (47.6% v:v), 10× PBS (1.35% v:v), rat tail collagen type I (8.2 mg/mL, Corning, 354249) (12.2% v:v), and 100 mM NaOH (2.8% v:v). The cell suspension was then added to the collagen mixture (36% v:v), yielding final concentrations of 1 mg/mL collagen and 12.6 × 10⁶ cells/mL. The mixture was gently mixed, and 10 μL or 50 μL was carefully dispensed into each well of a 15-well µ-Slide or a custom-made PDMS device, respectively. The dishes were placed on ice for 5 min and then transferred to a pre-heated microscope stage maintained at 37°C and 5% CO₂ to initiate collagen polymerization. After 7 min of polymerization, pre-warmed imaging medium was gently added to each well, and imaging was started immediately thereafter.

### Swarming and calcium dynamics imaging

Swarming experiments were imaged using IX-83 (Evident) inverted microscopes equipped with controlled atmosphere (37◦C, 5% CO2) using a ×20 dry objective (NA 0.80). To facilitate cell tracking at high cell density, dHL-60 cells were divided into three populations prior to embedding in collagen: one-third of the cells were labeled with Hoechst, one-third with SiR-DNA 647, and one-third were left unlabeled. The three populations were then mixed in equal proportions. Image acquisition was initiated immediately after the addition of imaging medium to the polymerized collagen gel. Fields of view were selected such that the zymosan spot was positioned at the center of the image. Bright-field, Hoechst, and SiR-DNA fluorescence images were acquired every 30 s throughout the experiment.

For comparison, 2D and 3D swarming experiments were performed in parallel using identical cell preparation and imaging settings in IBIDI 15-well µ-Slides. For 2D conditions, 7 × 10⁵ dHL-60 cells in 50 µL were directly added to wells containing pre-deposited zymosan spots at the same time the 3D samples were transferred to the microscope stage (37°C). Imaging was then initiated simultaneously for both conditions.

For intracellular calcium measurements, dHL-60 cells were loaded with 5 µM Fluo-8 AM (BioValley, XX) for 20 min prior to embedding in collagen gels. Alternatively, dHL-60 cells stably expressing the genetically encoded calcium indicator GCaMP6F were used. In both cases, fluorescence was acquired using 488 nm excitation and 525 nm emission every 20 s during swarming experiments. Where indicated, cells were pretreated with 2 mM EGTA for 15 min prior to collagen embedding to chelate extracellular calcium.

### HyPer7 imaging

Imaging was performed on a spinning-disk confocal microscope consisting of a Yokogawa CSU-X1 scan head mounted on a Leica DMi8 inverted microscope, equipped with a Hamamatsu ORCA-Flash4.0 camera, a NanoScanZ piezo focus drive (Prior Scientific), and a motorized XY stage (Marzhauser). The microscope was controlled using MetaMorph software (Molecular Devices) and fitted with an environmental chamber maintaining cells at 37°C and 5% CO₂ throughout imaging. For swarming experiments, images were acquired using a 20× objective. A single focal plane positioned near the glass surface, where most cells were located, was recorded every 20 s for 90 min. Ratiometric imaging of the HyPer7 probe was performed by sequential acquisition of two channels: an oxidized-state channel C2 (475 nm excitation, 525 nm emission) and a reduced-state channel C1 (405 nm excitation, 525 nm emission). Before each experiment, laser power and exposure times were adjusted to obtain comparable fluorescence intensities in both channels under basal conditions. A transmitted-light (bright-field) image was acquired at each time point. For experiments using fluorescently labeled zymosan particles (Zymosan A Alexa Fluor™ 594 conjugate, Thermo Fisher Scientific, Z23374), an additional fluorescence channel was acquired using 561 nm excitation.

For HyPer7 measurements following PMA or exogenous H_2_O_2_ stimulation, dHL-60 cells were pelleted by centrifugation and resuspended in Opti-MEM (Gibco, 31985-070) at a concentration of 2 × 10⁶ cells/mL. A volume of 100 µL of cell suspension was seeded into each well of a 96-well plate and incubated for 20–30 min at 37°C and 5% CO₂ to allow cell attachment. The medium was then gently aspirated and replaced with 100 µL of imaging medium. Baseline fluorescence was recorded for 10 min at a rate of one image per minute. Cells were subsequently stimulated by addition of 100 µL of imaging medium containing either PMA (phorbol 12-myristate 13-acetate; Thermo Fisher, CAS 16561-29-8) or H_2_O_2_ (Thermo Fisher, CAS 7722-84-1), yielding final concentrations of 5 or 100 nM PMA, or 20 or 200 µM H_2_O_2_, respectively. Imaging was then continued for an additional 45–60 min. Where indicated, cells were pretreated for 30 min before imaging with either 5 µM diphenyleneiodonium chloride (DPI; MedChemExpress, HY-100965) or 25 µM GSK2795039 (MedChemExpress, HY-18950).

### Image and Data Analysis

#### Ratiometric image processing

Raw image sequences were processed using Fiji/ImageJ (version 1.52p). The 405 nm (C1) and 488 nm (C2) channels were separated. A background subtraction was performed on each channel, followed by a mean filter (0.5 pixel radius). A binary mask was created from the C1 channel by applying a threshold to segment cells using pixel classification Ilastik or Labkit plugins. The mask was cleaned of small particles using Analyze Particles function and divided by 255. Both C1 and C2 were multiplied by this mask. Finally, a ratiometric image was generated by dividing the masked C2 image by the masked C1 image (HyPer7 Ratio = C2/C1).

#### Data extraction and normalization for HyPer7 during swarming

The processed image sequences were analyzed in Fiji. For each field of view, two regions of interest (ROIs) were defined. The “inside” region was manually drawn around the Zymosan spot area using the bright-field channel as a guide. The “outside” region was defined as the entire field of view excluding the inside region. The mean pixel intensity of the HyPer7 ratio signal was measured over time. To ensure comparability between different experiments, the data for each ROI was normalized to its own baseline by dividing each time point by the average intensity of the first five time points.

#### Tracking and trajectory analysis for swarming

Except for Figures 4A and 4B, where HyPer7 (C2 channel) was used for tracking, cell tracking was performed using fluorescently labeled nuclei. Images were preprocessed by background subtraction followed by application of a 1-pixel mean filter. The zymosan spot region was manually excluded from the analysis to remove cells located within the swarm core. Cell tracking was performed using the TrackMate plugin (Fiji), and tracks corresponding to cells that moved less than 40 µm over the entire imaging period (180 min) were excluded from the analysis. Cell positions (x, y coordinates) were extracted and further analyzed using custom-built software for trajectory analysis. This tool enables spatiotemporal analysis with selectable spatial and temporal windows. Radial directionality was defined as the velocity component directed toward the center of the zymosan spot divided by the total instantaneous speed.

### Statistical Analysis

Statistical analyses were performed using GraphPad Prism. Specific statistical tests applied are described in the corresponding figure legends. A p-value of less than 0.05 was considered statistically significant.

## SUPPLEMENTARY FIGURES

**Figure S1.**
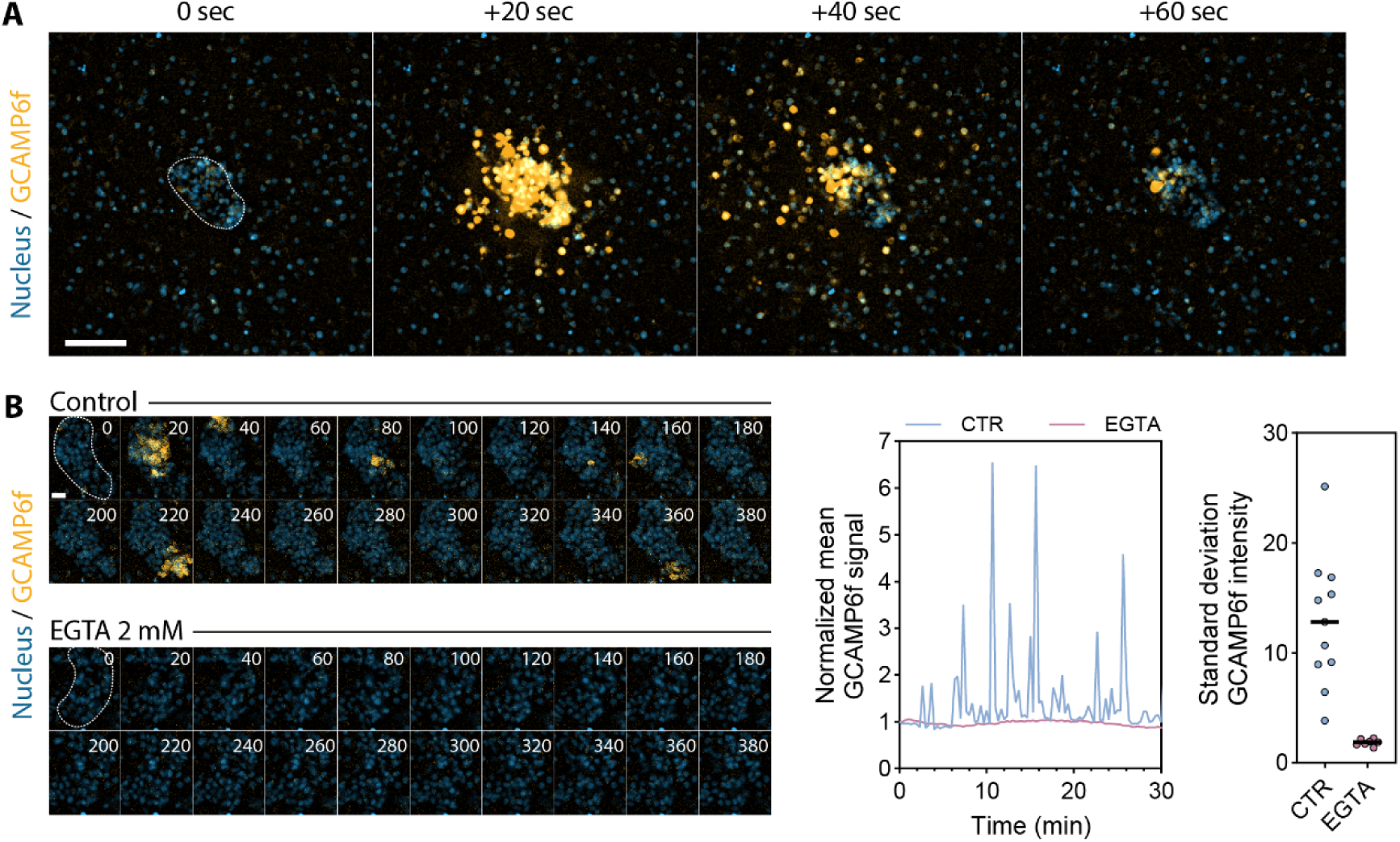
Calcium dynamics during swarming using GCaMP6F dHL-60. (A) Time-lapse sequence of a multicellular wave of neutrophil calcium activity following detection of the zymosan. The dotted line represents the zymosan spot boundary. Scale bar, 100 µm. (B) Left, time-lapse sequence of a calcium activity in the swarm core in the presence or not of 2 mM EGTA. The dotted line represents the zymosan spot boundary. Scale bar, 20 µm. Right, example the evolution of mean GCaMP6F signal in a swarm core (in the presence or not of EGTA) and standard deviation of GCaMP6F intensity. N = 8 swarms from n = 2 independent experiments.

**Figure S2.**
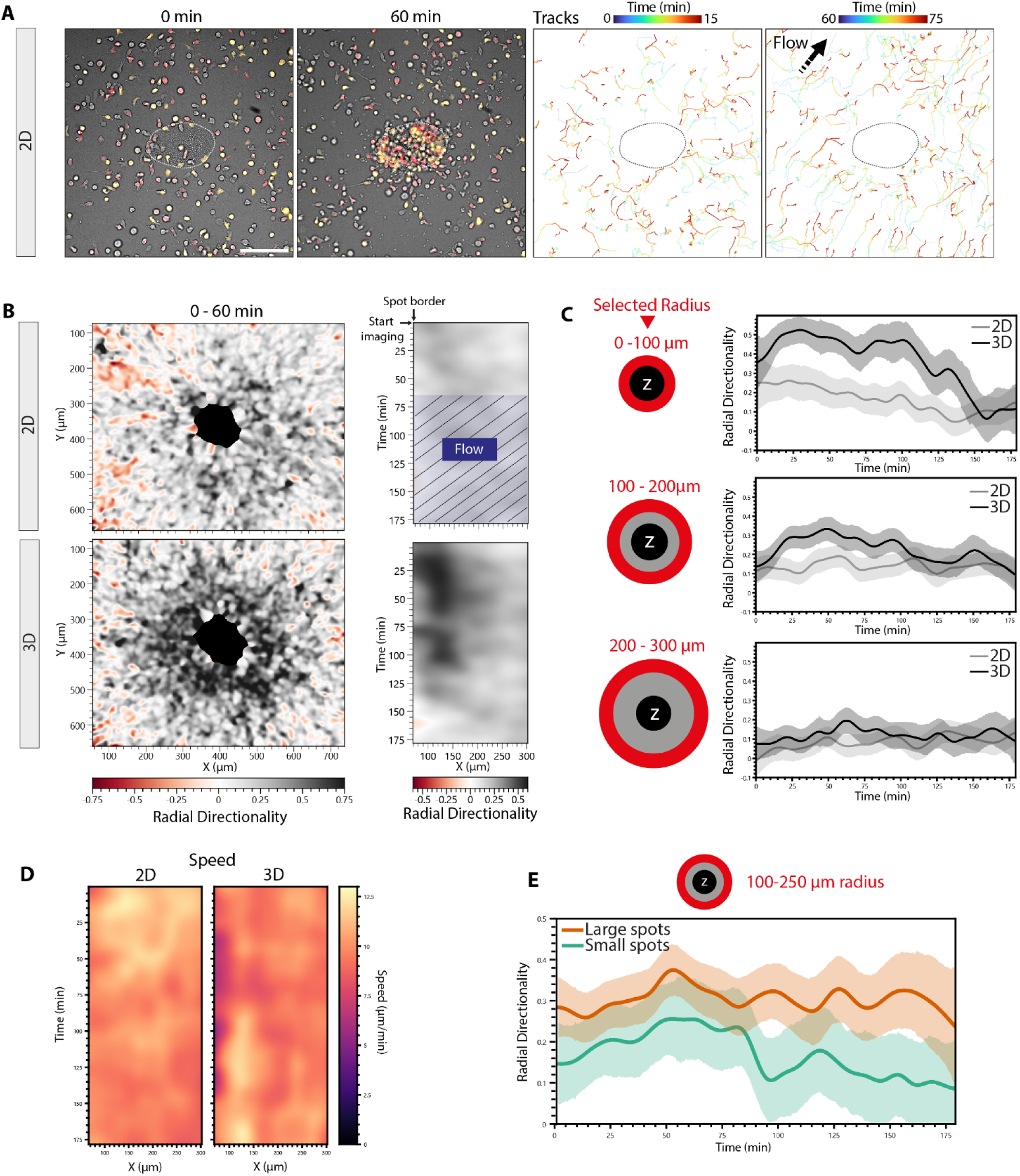
Comparison between 2D and 3D swarming of dHL-60. (A) Imaging and tracking of dHL-60 swarming in 2D. The dotted line represents the zymosan spot boundary. After 60 min, an important flow was observed. Scale bar, 100 µm. (B) Contour plot (XY at the left and XT at the right) showing mean radial directionality between 0 and 60 min of swarming imaging in 2D (top) and 3D (left). Data of 3D condition are the same than Figure 2C. N = 4 swarms from one representative experiment. (C) Evolution of mean radial directionality over time of dHL-60 in 2D and 3D depending on the region selected (red) based on the radius from the spot boundary. (D) Contour plot (XT) showing mean speed evolution over time of dHL-60 in 2D and 3D. N = 4 swarms from one representative experiment. (E) Evolution of mean radial directionality over time of dHL-60 in 3D depending on the size of zymosan spot.

**Figure S3.**
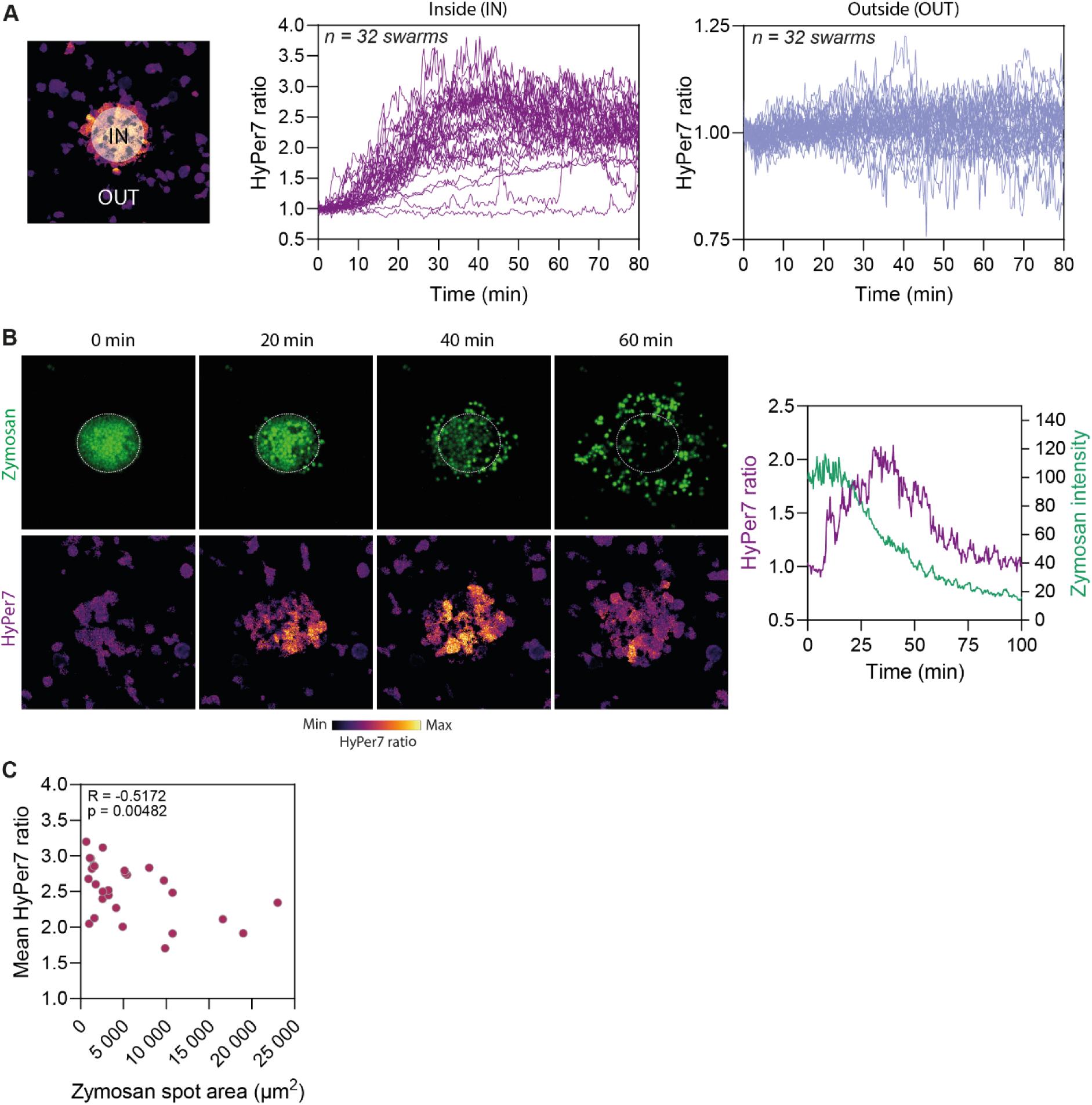
HL-60 HyPer7 cell line for studying spatiotemporal H_2_O_2_ production. (A) Individual swarm analysis of H_2_O_2_ levels inside and outside the initial zymosan area, during the first 80 min of imaging. N = 32 swarms from 5 independent experiments. Data corresponds to Figure 3D. (B) Left, imaging of the HyPer7 probe and fluorescent zymosan displacement during swarming. Right, quantification of HyPer7 ratio and zymosan intensity inside the initial zymosan area (dotted line region). (C) Correlation of mean HyPer7 ratio (between 30 and 40 min) with initial zymosan spot area. Spearman correlation coefficient r = –0.51. P < 0.01.

**Figure S4.**
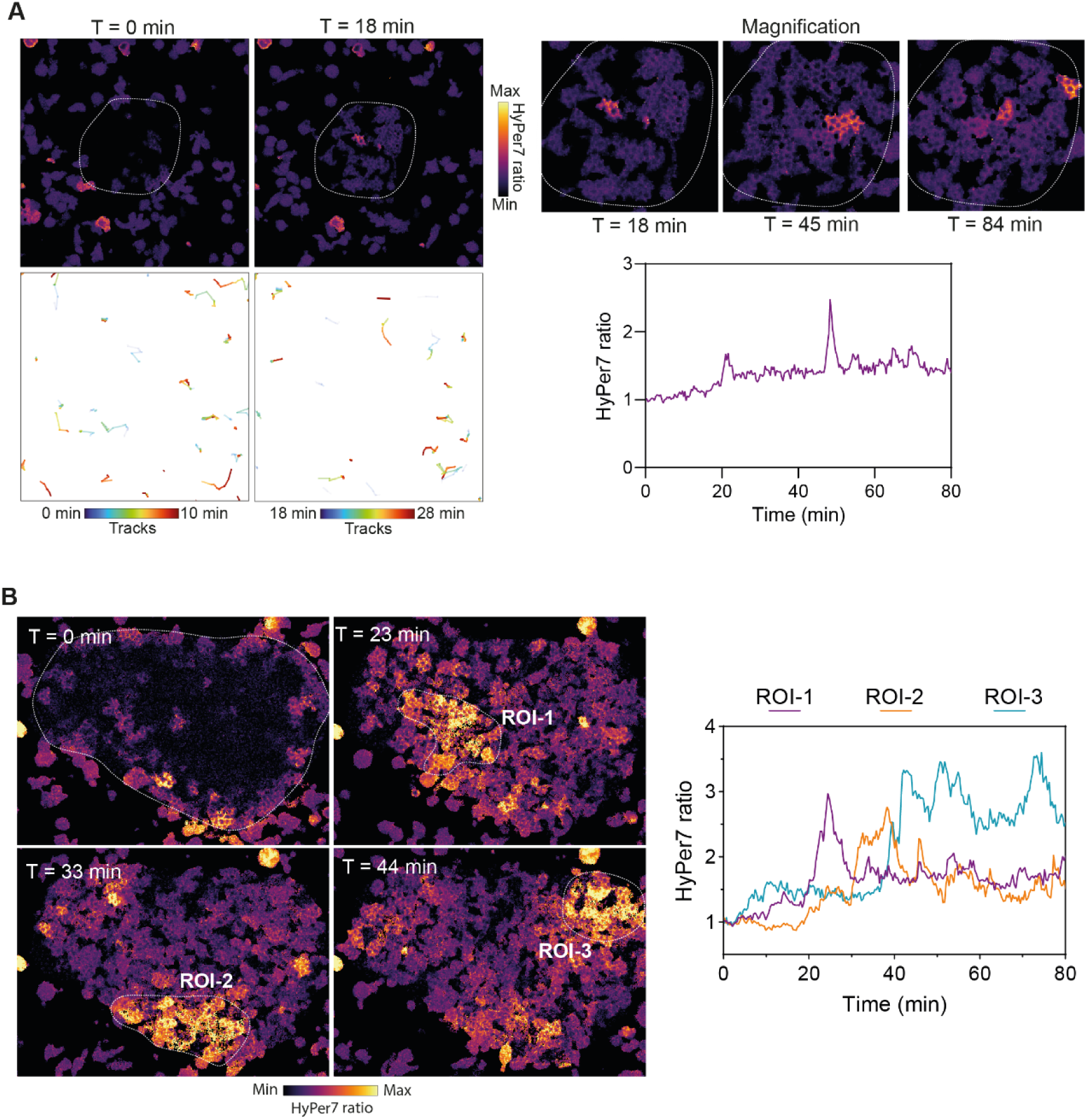
Examples of distinct H_2_O_2_ production behaviors. (A) Left, example showing no initiation of swarming and an absence of H**_2_**O**_2_** global production on zymosan area. The dotted line represents the zymosan spot boundary. Upper right, magnification of zymosan area showing transient intracellular H**_2_**O**_2_** production in individual cells. Lower right, quantification of mean HyPer7 ratio inside the zymosan area over 80 min. (B) Left, example of transient and local increase of intracellular H2O2 in subgroups of cells on a large zymosan spot. Right, quantification of mean HyPer7 ratio in three regions of interest (ROI) indicated with dotted lines on the images.

**Figure S5.**
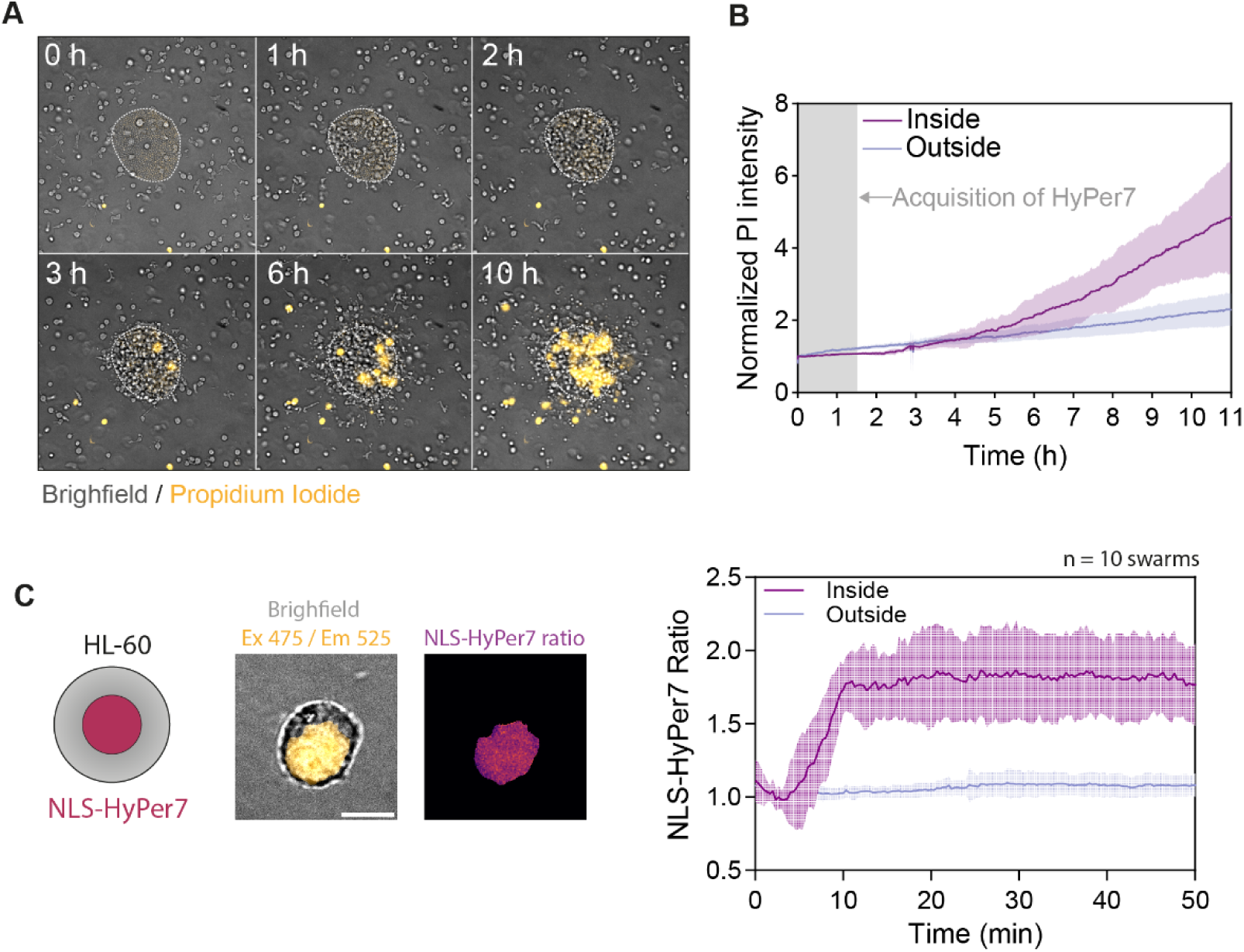
H_2_O_2_ production in the swarm, cell death and nuclear H_2_O_2_ localization. (A) Images of dHL-60 and cell permeabilization revealed by propidium iodide (PI). (B) Quantification of PI intensity showing that permeabilization of cells started from 3h after swarming initiation and increased specifically inside the zymosan area (swarm core). (C) Images of HL-60 expressing nuclear-targeted version of HyPer7 (NLS-HyPer7). Right, quantification of mean NLS-HyPer7 ratio inside and outside regions during dHL-60 swarming in 3D. N = 10 swarms from n = 2 independent experiments.

**Figure S6.**
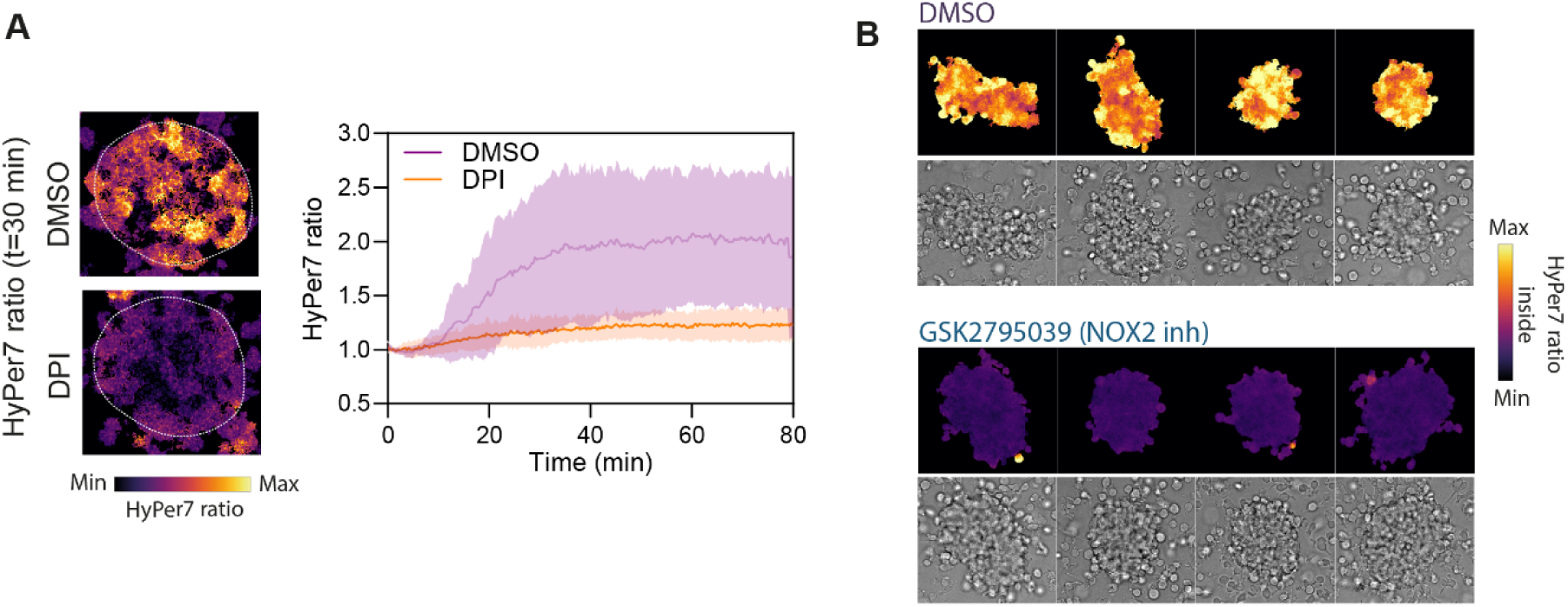
NOX2-dependent H_2_O_2_ production during swarming. (A) quantification of mean HyPer7 ratio (inside) for DMSO and DPI treated dHL-60. N = 17 (DMSO) and 20 (DPI) swarms from 3 independent experiments. (B) Representative images of HyPer7 ratio for 4 swarms of dHL-60 treated with DMSO or GSK2795039 after 1h of swarming. Images were taken with widefield microscope.

